# Deep Residual Network Reveals a Nested Hierarchy of Distributed Cortical Representation for Visual Categorization

**DOI:** 10.1101/151142

**Authors:** Haiguang Wen, Junxing Shi, Wei Chen, Zhongming Liu

## Abstract

The brain represents visual objects with topographic cortical patterns. To address how distributed visual representations enable object categorization, we established predictive encoding models based on a deep residual neural network, and trained them to predict cortical responses to natural movies. Using this predictive model, we mapped human cortical representations to 64,000 visual objects from 80 categories with high throughput and accuracy. Such representations covered both the ventral and dorsal pathways, reflected multiple levels of object features, and preserved semantic relationships between categories. In the entire visual cortex, object representations were modularly organized into three categories: biological objects, non-biological objects, and background scenes. In a finer scale specific to each module, object representations revealed sub-modules for further categorization. These findings suggest that increasingly more specific category is represented by cortical patterns in progressively finer spatial scales. Such a nested hierarchy may be a fundamental principle for the brain to categorize visual objects with various levels of specificity, and can be explained and differentiated by object features at different levels.

## Introduction

The visual cortex performs rapid categorization of complex and diverse visual patterns or objects. This ability is attributable to hierarchical neural computation and representation of category information^1,2^. In particular, the ventral temporal cortex contains topologically organized maps of object representations^3^, spanning a high-dimensional space^4^ while being invariant against changes in low-level visual properties^5,6^. Evidence shows that category representations also exist in the dorsal stream^7–9^ or even beyond^10^, likely reflecting non-visual attributes needed for category dependent actions^11,12^. It has thus been proposed that distributed cortical networks extract and represent category information for robust, efficient, and flexible visual categorization in multiple levels of abstraction^13,14^.

However, the computational understanding of distributed neural coding is still limited. Questions are unresolved as to how information is encoded in distributed patterns^14^, how object knowledge emerges from lower-level visual features^2^, and how cortical representations share and differ across categories^3,12^. Answering such questions requires a fully accessible computational model of hierarchical cortical processing for visual categorization^15^, and in principle to map the representations of as many categories as possible in a huge, if not infinite, dimension of object domains^16^. These challenges and requirements may be met by recent advances in deep neural networks (DNN)^17^ – a type of artificial neural networks built with conceptually similar architecture and computing principle as the brain itself ^2^.

Recent studies show that convolutional neural networks offer hierarchical representations of any visual input to be able to model and predict cortical responses to natural picture^18–22^ or video^23,24^ stimuli. The predictive power of such network models is high and robust in the entire visual cortex^24^, rendering them more effective than other models that only account for either the lowest^25,26^ or highest^16^ level in the visual hierarchy. The DNN-based predictive model can be applied to novel (unseen) visual stimuli^19,20,22,24^. It thus enables to simulate the cortical representations of a large number of visual objects and categories^22,24^, far beyond what is attainable experimentally^27–31^.

Extending from recent studies^18–22,24^, we used a deep residual network (ResNet)^32^ to define, train, and test a generalizable, predictive, and hierarchical model of natural vision by using extensive functional magnetic resonance imaging (fMRI) data from humans watching >10 hours of YouTube videos. Taking this predictive model as a “virtual” fMRI scanner, we synthesized the cortical response patterns with 64,000 natural pictures including objects from 80 categories, and mapped category representations in the human brain with high-throughput. We analyzed and compared the cortical representational similarity among categories against their semantic relationships, and quantitatively evaluated the differential contributions from different levels of visual features to the cortical organization of categories. Our results support the hypothesis that the brain uses nested spatial and representational hierarchies to perform multi-level visual categorization. Object representations in large to small scales support coarse to fine categorization^3^. In addition, different levels of categorization are primarily attributed to different levels of object knowledge, with greater contributions from middle and high-level object features than from low-level image features.

## Results

### ResNet predicted widespread cortical responses to natural visual stimuli

In line with recent studies^18–22,24^, we used a deep convolutional neural network to establish predictive models of the cortical fMRI representations of natural visual stimuli. Specifically, we used the ResNet – a deep residual network pre-trained for computer vision^32^, with a much deeper architecture to yield more fine-grained layers of visual features than otherwise similar but shallower networks, e.g. AlexNet^33^ explored in prior studies^18–22,24,34^. From any visual stimuli, the ResNet-extracted features jointly predicted the fMRI response through a voxel-wise linear regression model. This encoding model was trained with a large amount of fMRI data during a training movie (12.8 hours for Subject 1, and 2.4 hours for Subject 2, 3), and tested with an independent testing movie (40 minutes).

The encoding accuracy (i.e. the correlation between the predicted and measured fMRI signals during the testing movie) was overall high (r = 0.43*±*0.14, 0.36*±*0.12, and 0.37*±*0.11 for Subject 1, 2 and 3, respectively) and statistically significant (permutation test, corrected at FDR q<0.01) throughout the visual cortex in every subject (Fig. 1.a). The encoding accuracy was comparable among the higher-order ventral-stream areas, e.g. fusiform face area (FFA) and parahippocampal place area (PPA), as well as early visual areas, e.g. V1, V2, V3, and V4 (Fig. 1.c), whereas it was relatively lower at such dorsal-stream areas as lateral intraparietal area (LIP), frontal eye fields (FEF), parietal eye fields (PEF), but not the middle temporal area (MT) (Fig. 1.c). Different cortical regions were found to be preferentially correlated with distinct layers in ResNet. The lower to higher level visual features encoded in ResNet were gradually mapped onto areas from the striate to extrastriate cortex along both ventral and dorsal streams (Fig. 1.b), in agreement with previous studies^20–24,34,35^. The prediction accuracy was consistently higher with (the deeper) ResNet than with (the shallower) AlexNet (Fig. 1.c). These results suggest that the ResNet-based voxel-wise encoding models offer generalizable computational accounts for the complex and nonlinear relationships between natural visual stimuli and cortical responses at widespread areas involved in various levels of visual processing.

**Figure1.**
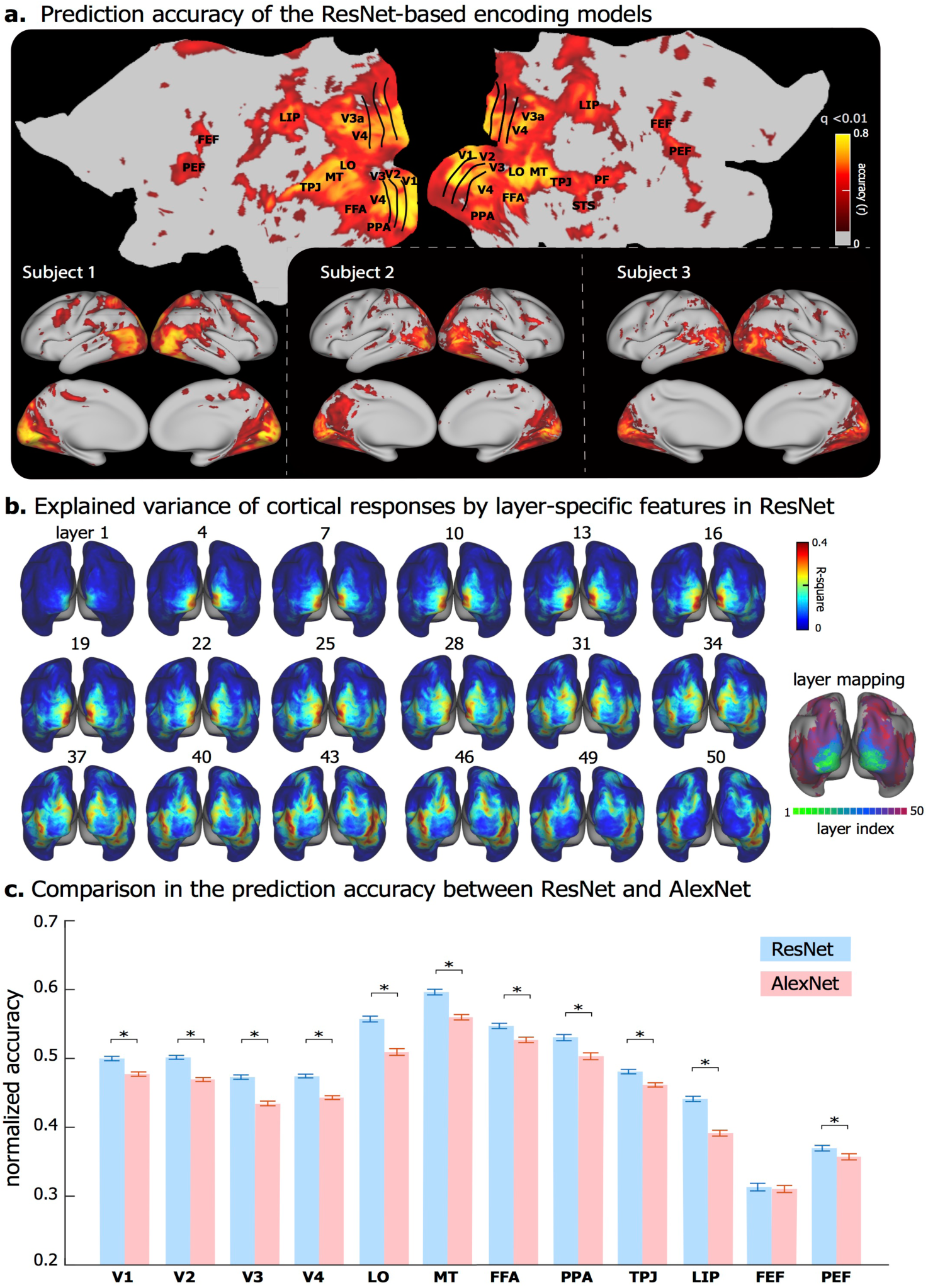
DNN-based Voxel-wise encoding models. **(a)** Performance of ResNet-based encoding models in predicting the cortical responses to novel testing movies for three subjects. The accuracy is measured by the average Pearson’s correlation coefficient (r) between the predicted and the observed fMRI responses across five testing movies (q<0.01 after correction for multiple testing using the false discovery rate (FDR) method, and with threshold r>0.2). The prediction accuracy is displayed on both flat (top) and inflated (bottom left) cortical surfaces for Subject 1. **(b)** Explained variance of the cortical response to testing movie by the layer-specific visual features in ResNet. The right shows the index to the ResNet layer that most explains the cortical response at every voxel. **(c)** Comparison between the ResNet-based and the AlexNet-based encoding models. Each bar represents the mean*±*SE of the prediction accuracy (normalized by the noise ceiling, i.e. dividing prediction accuracy (r) by the noise ceiling at every voxel) within a ROI across voxels and subjects, and * represents a significance p-value (p<0.001) with paired t-test.

### Encoding models predicted cortical representations of various object categories

As explored before ^22,24^, the voxel-wise encoding models constituted a high-throughput computational workbench to synthesize cortical activations with a very large number of natural pictures, not realistically attainable with most experimental approaches. Here, we used this strategy to predict the pattern of cortical activation with each of the 64,000 natural pictures from 80 categories with 800 exemplars per category. By averaging the predicted activation maps across all exemplars of each category, the common cortical activation within this category was obtained to report its cortical representation.

For example, averaging the predicted responses to various human faces revealed the category-wide cortical representation of the “face” invariant of low-level visual features, e.g. the color, position, and perspective (Fig. 2.a). Such a model-simulated “face” representation was consistent with the fMRI-mapping result obtained with a block-design functional localizer that contrasted face vs. non-face pictures (Fig. 2.b). In a similar manner, cortical representations of all 80 categories were mapped (Fig. 3). The resulting category representations were not only along the ventral stream, but also along the dorsal stream albeit with relatively lower amplitudes and a smaller extent.

**Figure2.**
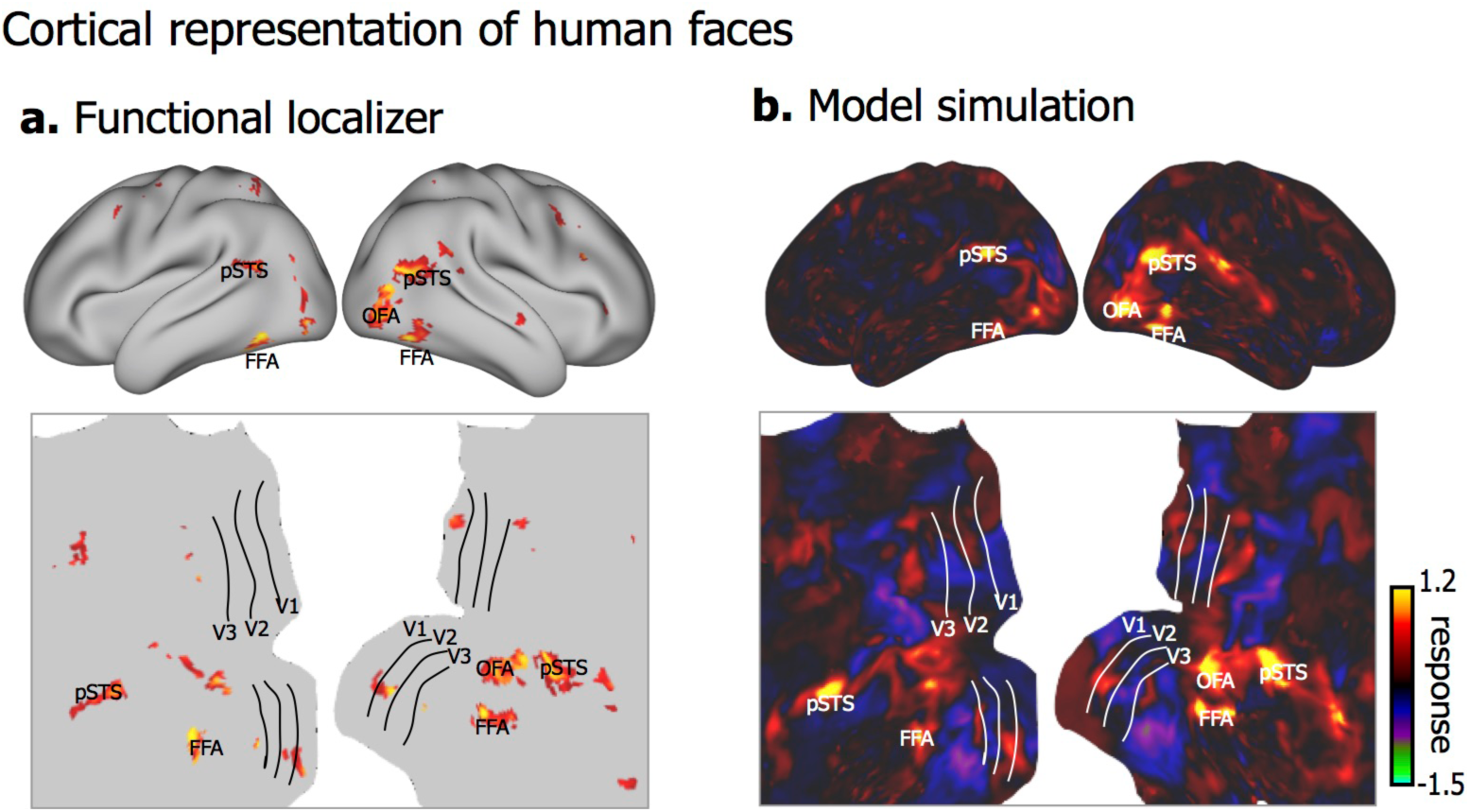
Human-face representations with encoding models and functional localizer. **(a)** Model-simulated representation of human face from ResNet-based encoding models. The representation is displayed on both inflated (top) and flat (bottom) cortical surfaces. **(b)** Localizer activation maps comprising regions selective for human faces, including occipital face area (OFA), fusiform face area (FFA), and posterior superior temporal sulcus (pSTS).

**Figure3.**
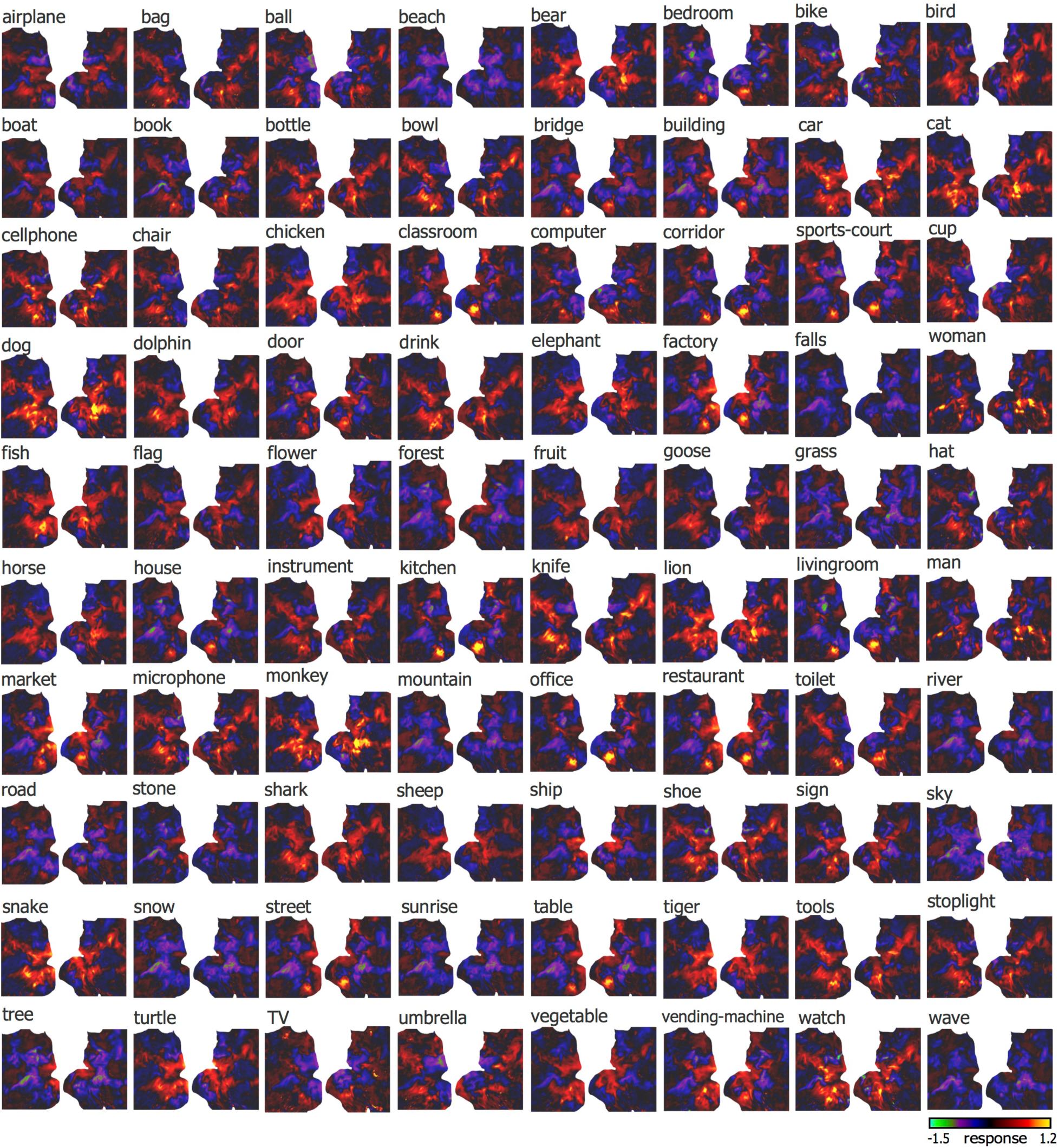
Cortical representations of 80 object categories. Each panel shows the representation map of an object category on flat cortical surface from Subject 1. The category label is on top left. The color bar shows the cortical response.

For each voxel, the model-predicted response as a function of category was regarded as the voxel-wise profile of categorical representation. The category selectivity – a measure of how a voxel was selectively responsive to one category relative to others ^36^, varied considerably across cortical locations (Fig. 4.a). Voxels with higher category selectivity were clustered into discrete regions including the bilateral PPA, FFA, lateral occipital (LO) area, the temporo-parietal junction (TPJ), as well as the right superior temporal sulcus (STS) (Fig. 4.a). The profile of categorical representation listed in a descending order (Fig. 4.b), showed that FFA, OFA, and pSTS were selective to humans or animals (e.g. man, woman, monkey, cat, lion); PPA was highly selective to places (e.g. kitchen, office, living room, corridor); the ventral visual complex (VVC) was selective to man-made objects (e.g. cellphone, tool, bowl, car). In general, the ventral stream tended to be more category-selective than early visual areas (e.g. V1, V2, V3) and dorsal-stream areas (e.g. MT, LIP) (Fig. 4.c).

**Figure4.**
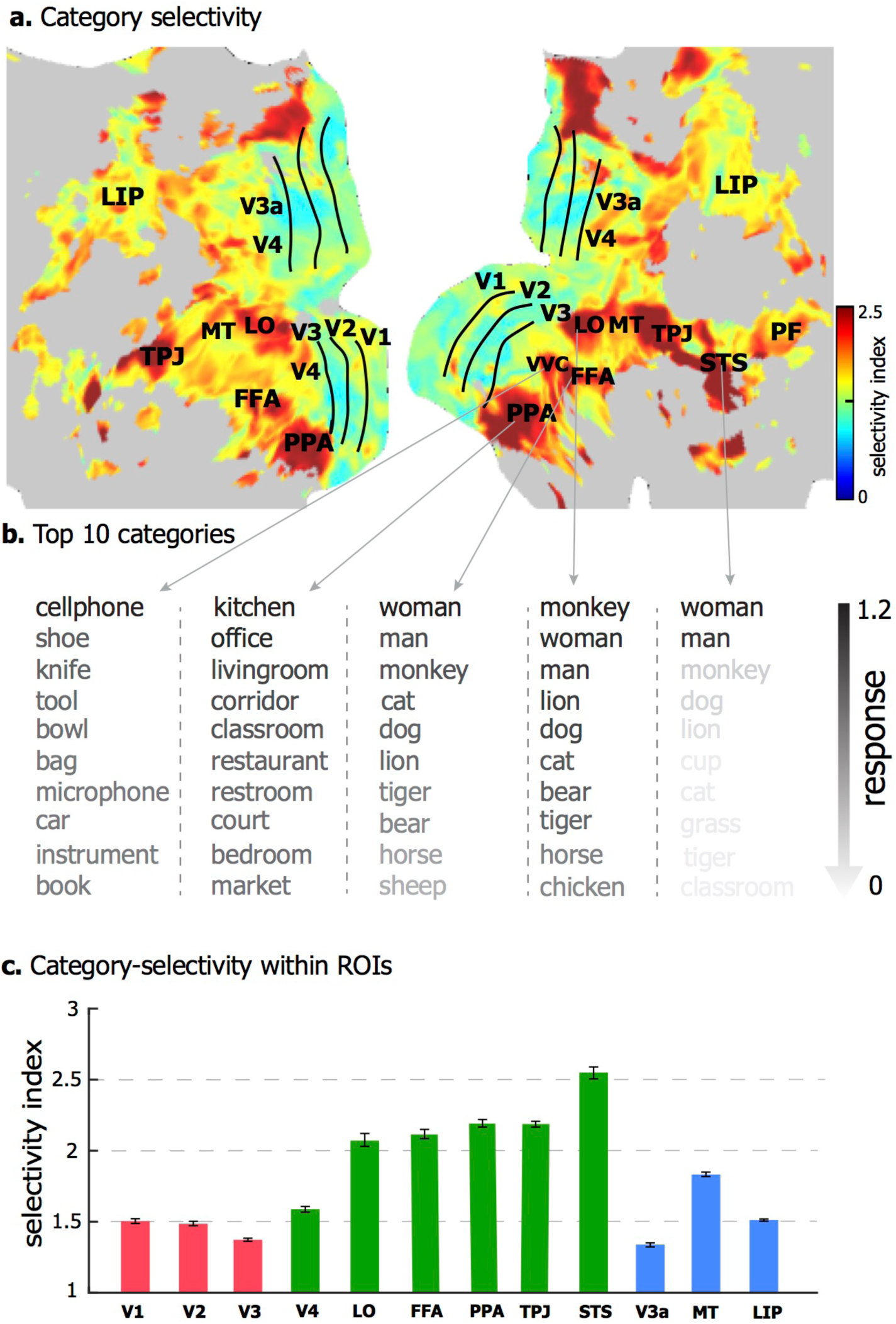
Category-selectivity at individual cortical locations. **(a)** The category-selectivity across the cortical surface. **(b)** The category-selectivity profile of example cortical locations. For each location, top 10 categories with the highest responses are showed in descending order. **(c)** Category-selectivity within ROIs (mean*±*SE) in the early visual areas (red), ventral stream areas (green), and dorsal stream areas (blue).

### Distributed, overlapping, and modular representations of categories

Although some ventral-stream areas (e.g. PPA and FFA) were highly (but not exclusively) selective to a certain category, no category was represented by any single region alone (Fig. 3). As suggested previously^14^, object categories were represented distinctly by distributed but partially overlapping networks (see examples in Supplementary Fig. S1 online). In the scale of the nearly entire visual cortex as predictable by the encoding models (Fig. 1.a), the spatial correlations in cortical representation between distinct categories were shown as a representational similarity matrix (Fig 5.a). This matrix revealed a modular organization (modularity Q=0.35), by which categories were clustered into three superordinate-level modules (Fig. 5.a, left). The categories being clustered based on their cortical representations exhibited a similarly modular pattern in terms of their semantic similarity (Fig. 5.a, middle), measured as the LCH similarity between the corresponding labels in WordNet ^37^. Interestingly, the similarity in cortical representation between categories was highly correlated with their semantic similarity (Fig. 5.a, right), suggesting that categories with more similar cortical representations tend to bear more closely related semantic meanings.

**Figure5.**
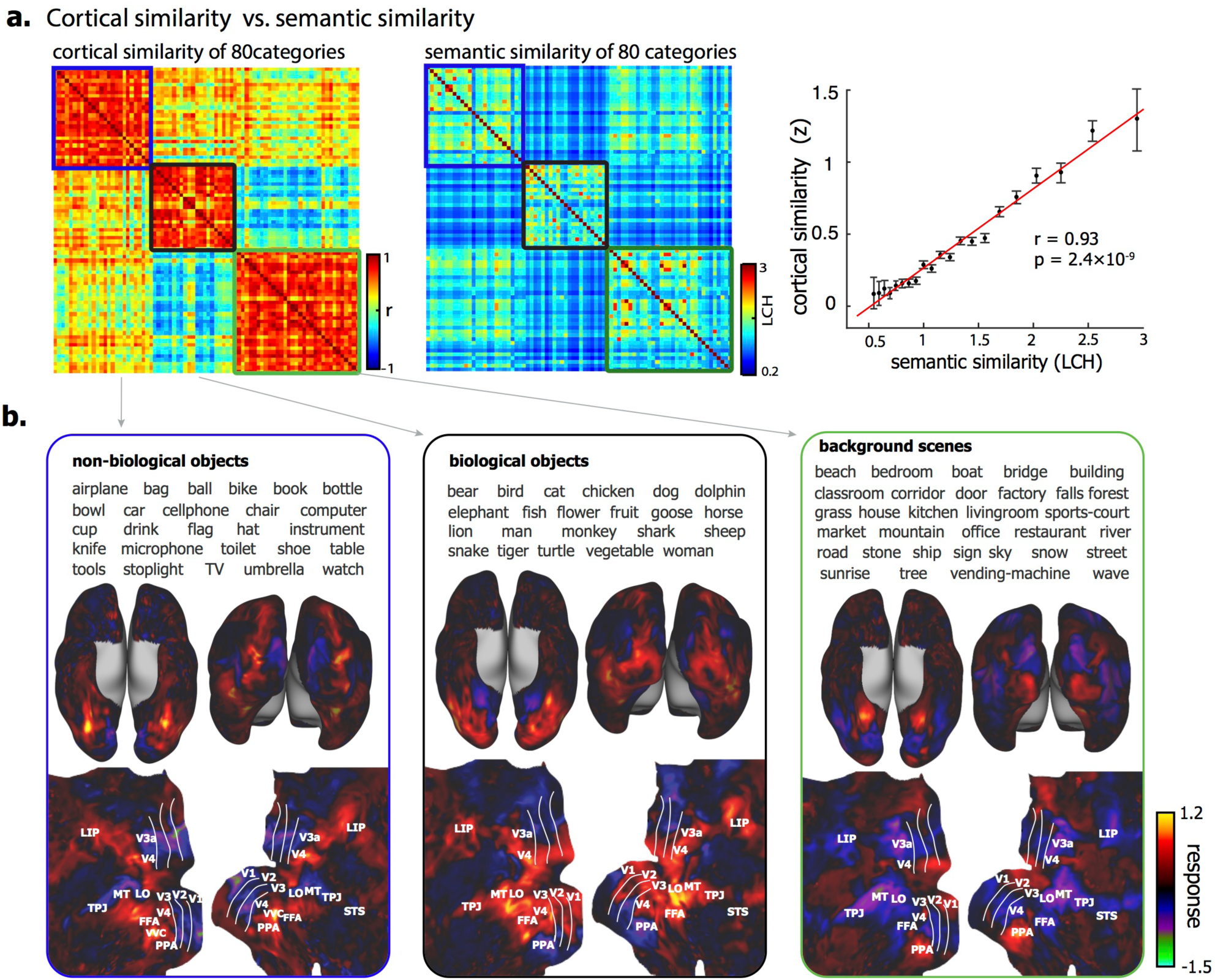
Categorical similarity and modularity in cortical representation at the scale of the entire visual cortex. **(a)** The left is the similarity matrix (r) of the cortical representations between categories. Each element represents the average cortical similarity between a pair of categories across subjects (see individual results in Supplementary Fig. S2 online). It is well separated into three modules with modularity Q=0.35. The middle is the similarity matrix of the semantic content between categories (measured by LCH). The right is a plot of the mean±SE of cortical similarity (Fisher’s z-transformation of r) vs. the semantic similarity (LCH takes discrete values). **(b)** These three modules are related to three superordinate-level categories: non-biological objects, biological objects, and background scenes. The average cortical representations across categories within modules are showed in the bottom on both inflated and flat cortical surfaces.

The representational modules in the entire visual cortex revealed coarse categories that seemed reasonable. The first module included non-biological objects, e.g. airplane, bottle and chair; the second module included biological objects, e.g. humans, animals, and plants; the third module included places and scenes (Fig. 5.b). The cortical representation averaged within each module revealed the general cortical representations of non-biological and biological objects, and background scenes (Fig. 5.b). As shown in Fig. 5.b, non-biological objects were represented by activations in bilateral sub-regions of ventral temporo-occipital cortex (e.g. VVC); biological objects were represented by activations in the lateral occipital cortex and part of the inferior temporal cortex (e.g. FFA) but deactivations in parahippocampal cortex (e.g. PPA); background scenes were represented by activations in PPA but deactivations in the lateral occipital complex, partly anti-correlated with the activations with biological objects.

### Mid-level visual features primarily accounted for basic-level categorization

Which levels of visual features accounted for such a modular organization were revealed by examining the representational similarity and modularity as attributed to the features extracted by each layer in the ResNet. Fig. 6.a (left) shows the inter-category representational similarity given the layer-wise features, thus decomposing the modular organization in Fig. 5.a by layers. The layer-wise modularity in cortical representation emerged progressively, being the lowest for the 1^st^ layer, showing noticeable three modules from the 10^th^ layer, and reaching the maximum at the 31^st^ layer (Fig. 6.a).

**Figure6.**
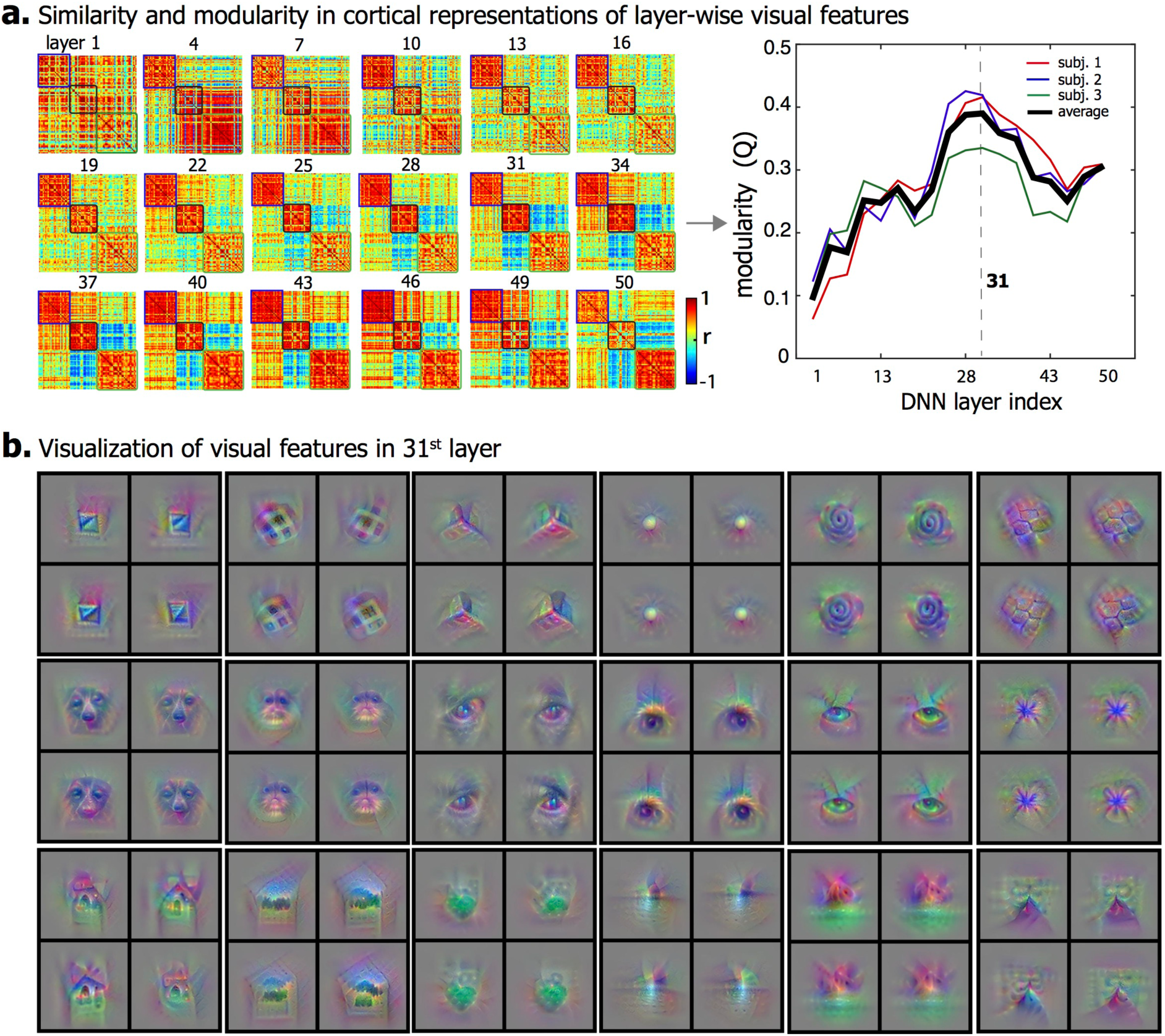
Contribution of layer-wise visual features to the similarity and modularity in cortical representation. **(a)**The left shows the similarity between categories in the cortical representations that are contributed by separated category information from individual layers. The order of categories is the same as in Figure 6.a. The right plot shows the modularity index across all layers. The visual features at the middle layers have the highest modularity. **(b)** 18 example visual features at the 31^st^ layer are visualized in pixel space. Each visual feature shows 4 exemplars that maximize the feature representation.

To gain intuition about the types of visual information from the 31^st^ layer, the features encoded by individual units in this layer were visualized. Fig. 6.b illustrates the visualizations of some example features, showing shapes or patterns (both 2-D and 3-D), animal or facial parts (e.g. head and eye), environmental components (e.g. house and mountain). Beyond these examples, other features were of similar types. Therefore, the mid-level features that depict object shapes or parts are modularly organized by their distributed cortical representations, supporting the superordinate-level categorization.

### More specific categories were modularly organized in finer scales

We further asked whether the similar modular organization could be extended to a lower level of categorization. That is, whether object representations were modularly organized within each superordinate-level module. For this purpose, we confined the scope of analysis from the whole visual cortex (Fig. 1.a) to finer spatial scales highlighted by co-activation patterns within biological objects, non-biological objects, or background scenes (Fig. 7.a). For example, within the regions where biological objects were represented (Fig. 7.a, top), the representational patterns were further clustered into four sub-modules: terrestrial animals, aquatic animals, plants, and humans (Fig. 7.b, top). Similarly, the fine-scale representational patterns of background scenes were clustered into two sub-modules corresponding to artificial (e.g. bedroom, bridge, restaurant) and natural scenes (e.g. falls, forest, beach) (Fig. 7, middle). However, non-biological objects showed a much less degree of modularity in cortical representation; the two modules did not bear any reasonable conceptual distinction (Fig. 7, bottom).

**Figure7.**
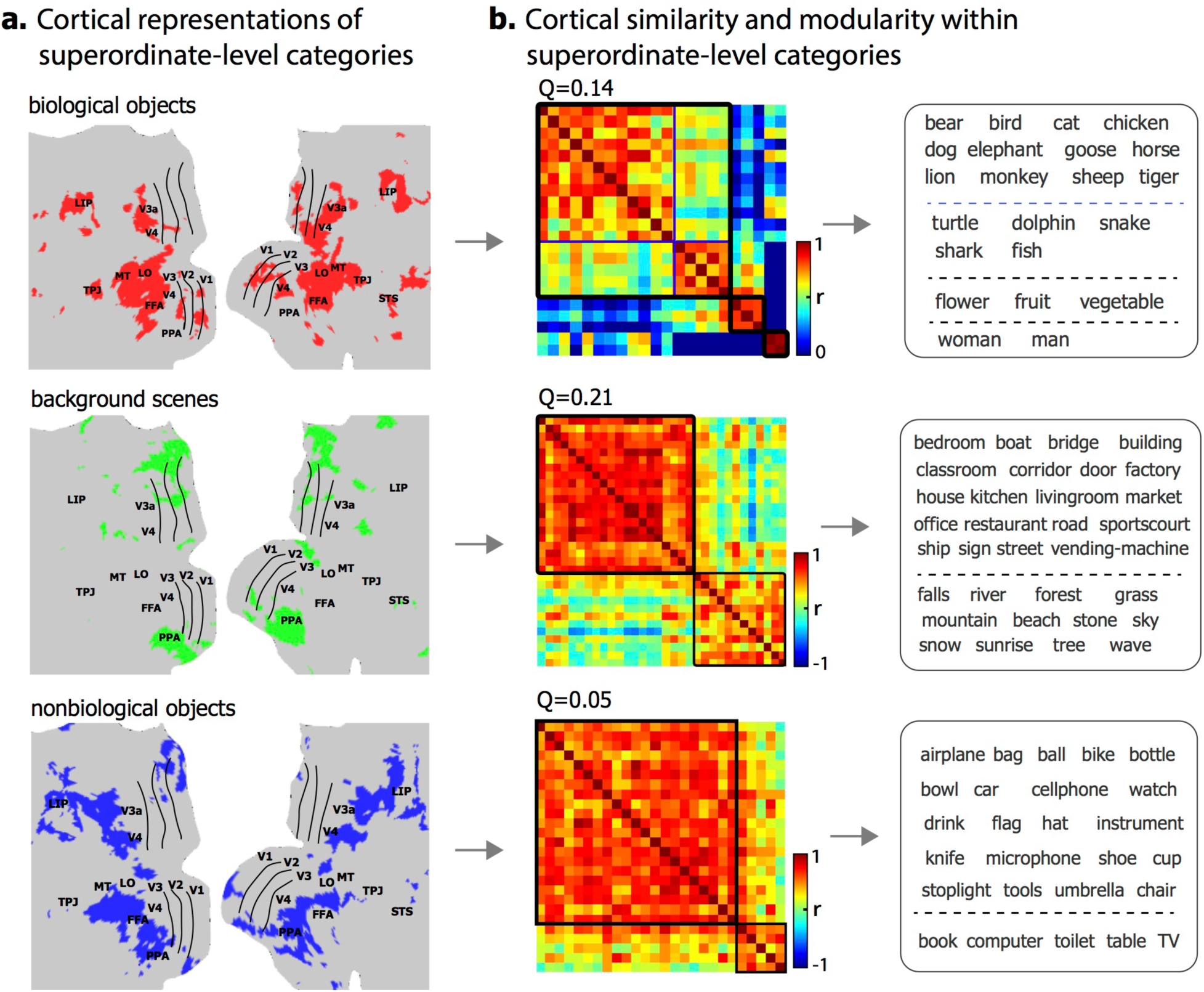
Categorical similarity and modularity in cortical representation within superordinate-level categories. **(a)** Fine-scale cortical areas specific to each superordinate-level category: biological objects (red), background scenes (green) and non-biological objects (blue). **(b)** The cortical similarity between categories in fine-scale cortical representation. The categories in each sub-module were displayed on the right. See individual results in Supplementary Fig. S2 online.

We also evaluated the layer-wise contribution of visual features to the fine-scale representational similarity and modularity. For biological objects, the modularity index generally increased from the lower to higher layer, reaching the maximum at the highest layer (Fig. 8.a, top). Note that the highest layer encoded the most abstract and semantically relevant features, whose visualizations revealed the entire objects or scenes (Fig. 8.b) rather than object or scenic parts (Fig. 6.b). In contrast, the modularity index reached the maximum at the 28^th^ layer for background scenes (Fig. 8.a, middle), but was relatively weak and less layer-dependent for non-biological objects (Fig. 8.a, bottom).

**Figure8.**
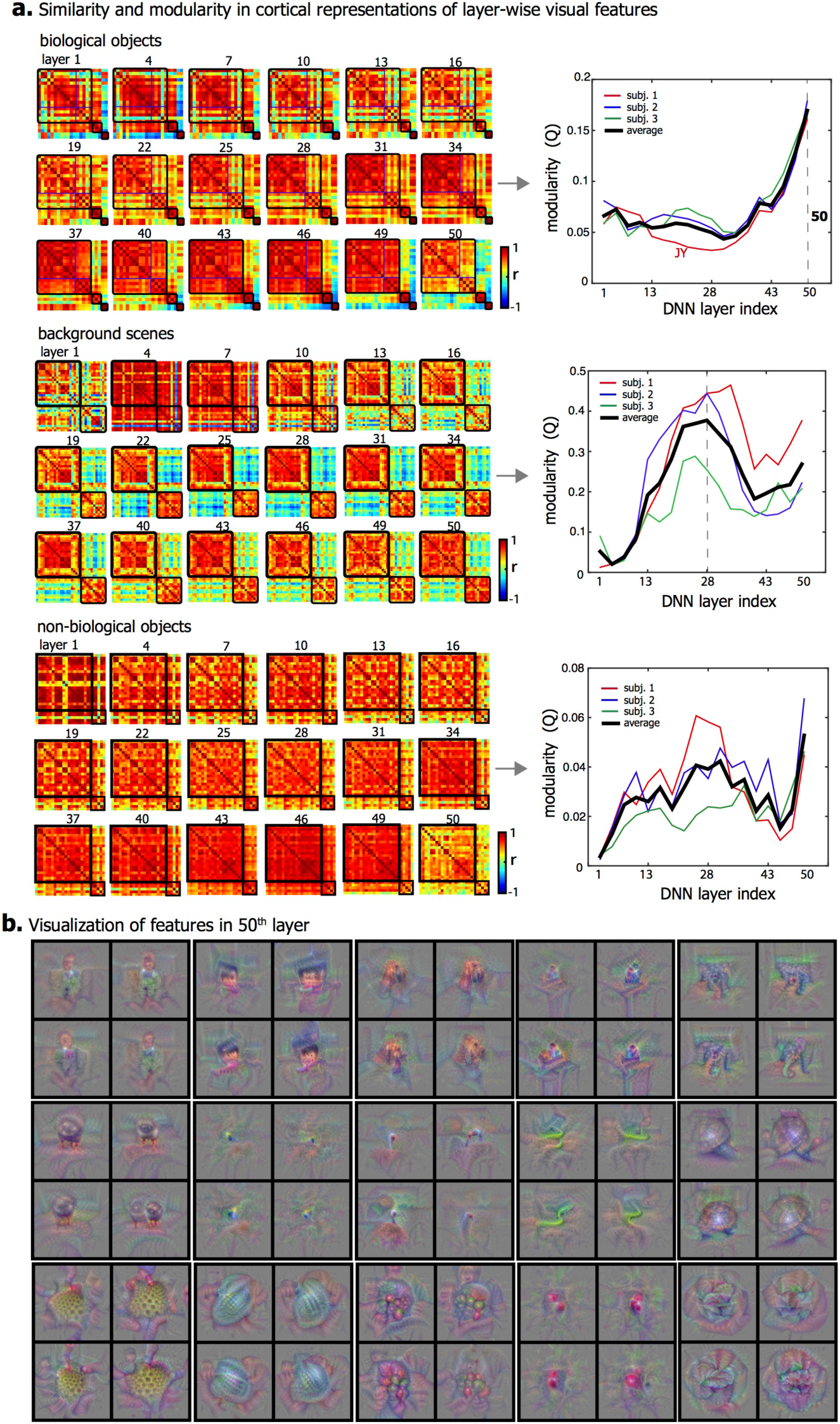
Contribution of layer-wise visual features to the similarity and modularity in cortical representations within superordinate-level categories. **(a)** The left shows the similarity between categories in fine-scale cortical representations that are contributed by separated category information from individual layers. The order of categories is the same as in Figure 8. The right plot shows the modularity index across all layers. The highest-layer visual features show the highest modularity for biological objects. **(b)** 15 example visual features at the 50^st^ layer are visualized in pixel space. Each visual feature showed 4 exemplars that maximize the feature representation.

## Discussion

This study demonstrates a high-throughput computational strategy to characterize hierarchical, distributed, and overlapping cortical representations of visual objects and categories. Results suggest that information about visual-object category entails multiple levels and domains of features represented by distributed cortical patterns in both ventral and dorsal pathways. Categories with similar cortical representations are more related in semantics. In a large scale of the entire visual cortex, object representations are modularly organized into three superordinate categories (biological objects, non-biological objects, and background scenes). In a finer scale specific to each module, category representation reveals sub-modules for finer categorization (e.g. biological objects are categorized into terrestrial animals, aquatic animals, plants, and humans). These findings support a nested hierarchy in distributed cortical representation for visual categorization: increasingly more specific category information is represented by distinct cortical patterns in progressively finer spatial scales ^3^, enabling the brain to identify, relate, and separate objects in various levels of abstraction. Meanwhile, the nested cortical organization of categories is primarily driven by object features from middle to high levels, rather than low-level image features.

Central to this study is the use of the categorization-driven deep ResNet for synthesizing the cortical representations of thousands of natural visual objects from many categories. This strategy has a much higher throughput for sampling a virtually infinite object or category space^22,24^, compared to prior studies that are limited to fewer categories with much fewer exemplars per category^27–31^. The sample size could be further extendable, since the ResNet-based encoding models account for the relationships between cortical responses and hierarchical and invariant visual features. Such features are finite and generalizable to different and new natural images, objects, and categories which the models have not been explicitly trained with. The model predictions are highly accurate and consistent with experimentally observed cortical responses (Fig. 1.a) and object representations (Fig. 2). The encoding accuracy may be further improved given an even larger and more diverse video-fMRI dataset to train the model, and a more biologically relevant deep neural net that better matches the brain and better performs in computer-vision tasks^19^. In this sense, the encoding models in this study are based on so far largest video-fMRI training data from single subjects; and ResNet also outperforms AlexNet in categorizing images^32,33^ and predicting the brain (Fig. 1.c). The encoding models reported here are thus arguably more powerful in predicting and mapping hierarchical cortical representations across the entire visual cortex (Fig. 1), compared to conceptually similar models in prior studies^18–22,24^.

What is also advantageous is that ResNet decomposes category information into multiple layers of features progressively emerging from low to mid to high levels. As such, ResNet offers a computational account of hierarchical cortical processing for categorization, yielding quantitative description of every object or category in terms of different layers of visual features. Mapping the layer-wise features from the ResNet onto the brain helps to address what drives the cortical organization of object knowledge and supports various levels of categorization.

The ResNet is trained with large-scale image set (~1.3 million natural images) for recognizing 1,000 visual object categories^32^. Though specific categories are used in training the ResNet, the trained model is generalizable to represent the semantics in our training and testing stimuli (Fig. 1), and is transferrable for recognizing new categories based on the generic representations in the learned feature space for transfer learning^38,39^. The generalizability of the feature space allows to predict the cortical representations of a wide range of categories far beyond those that the network has been explicitly trained. For example, the model is able to predict the face representation even though the ResNet is not trained for recognizing faces (Fig. 2).

Our results support the notion that visual-object categories are represented by distributed and overlapping cortical patterns ^14^ rather than clustered regions^40–42^. Given this notion, the brain represents a category not as a single entity but a set of defining attributes that span multiple domains and levels of object knowledge. Different objects may bear overlapping representational patterns that are both separable and associable, allowing them to be recognized as one category in a particular level, but as different categories in another level. For example, a lion and a shark are both animals but can be more specifically categorized as terrestrial and aquatic animals, respectively. The distributed and overlapping object representations, as weighted spatial patterns of attribute-based representations^11^, constitute an essential principle underlying the brain’s capacity for multi-level categorization.

Category representations, although distributed in general, may become highly selective at spatially clustered regions^40–42^. The category-selective regions are mostly in the ventral temporal cortex (Fig. 4), e.g. the FFA, PPA, and LO. The existence of category-selective regions does not contradict with distributed category representation. Instead, the category specificity in a region is thought to emerge from its connectivity with other regions that also represent that category^43^, for processing domain-specific knowledge of particular importance to vision-guided action and cognition ^44^.

The cortical representational similarity between different categories is highly correlated with their semantic relationship (Fig. 5). In other words, the semantic relationship is preserved by cortical representation. This finding lends support for the notion of a continuous semantic space underlying the brain’s category representation^16^, which is a compelling hypothetical principle to bridge neural representation and linguistic taxonomy^45^. However, category information is not limited to semantic features, but includes hierarchically organized attributes that all define categories and their conceptual relationships. For example, “face” is not an isolated concept; it entails facial features (“eyes”, “nose”, “mouth”), each also having its own defining features. The similarity and distinction between categories may be attributable to one or multiple levels of features. In prior studies^16^, the hierarchical nature of category information is not considered as every exemplar of each category is annotated by a pre-defined label. This causes an incomplete account of category representation, leaving it difficult to pinpoint what dimensions of category information drive the representational similarity between categories.

We have overcome this limit by extracting multiple layers of features from visual objects and evaluating the layer-wise contributions to cortical category representation. Our results show that similarity in cortical category representation is contributed by multiple layers of visual features, while different layers contributed differently to the representational modularity. Coarse categories (i.e. biological objects, non-biological objects, and background scenes) are most attributable to mid-level features, e.g. shapes, textures, and object parts (Fig. 6). In a finer level of categorization, terrestrial animals, aquatic animals, plants, and humans are most distinguishable in the semantic space; categorization of man-made and natural scenes is most supported by mid-level features (Fig. 8), likely reflecting the spatial layout of scene components^46–48^.

Our results suggest that object representations in the entire visual cortex support coarse categorization (Fig. 5), and representations in a smaller scale specific to each coarse category support subsequently finer categorization (Fig. 7). This finding is in line with the notion of nested spatial and representational hierarchies^3^: increasingly specific categorization results from category representations in a progressively finer spatial scale on the cortex. Such a spatial hierarchy describes a functional architecture that complies with both distributed^12,14^ and regional^40,42^ representations of object knowledge, and their functional roles for categorization in multiple levels of specificity^3^. This nested hierarchy implies that widely distributed patterns of responses to visual objects are more distinguishable between coarsely defined categories than between relatively finer categories, as demonstrated in previous studies^27,49^. Finer object categorization may require representational differences in domain-specific regions^40–42^.

The notion of spatial and representational hierarchies for graded categorization also has implications to decoding visual objects with multi-voxel pattern analysis^14,50–55^. No single spatial scale is optimal for decoding visual objects across all levels of categories. The optimal spatial scale for decoding object categories depends on how specific the categories are defined.

One of the unresolved questions about object categorization is about what dimensions drive the organization of categories in the brain^3,43,56^. A number of studies have suggested that the cortical representation in the ventral visual pathway is highly related to the categorical or semantic information of visual objects^31,49,57–59^, which is also shown in our results (Fig. 5a). Recent studies also suggest that the cortical organization of object categories can be explained by variance in low-level visual features^60–62^, shape similarity^61,63–67^, and the real-word or conceptual size of objects^68,69^. However, these known dimensions only partially explain the cortical organization, and it is currently unclear whether other dimensions, e.g. mid-level visual features ^70^, might drive the organization^3,43,56^. Thus, it is more desirable to evaluate and compare a more complete set of visual dimensions in explaining the categorical organization.

In this study, we used a much larger set of visual dimensions defined in ResNet^32^, including low (e.g. edges), mid (e.g. object parts), and high-level (e.g. semantic meaning) visual features^71^, to investigate the cortical organization of 64,000 visual objects over 80 categories. By quantitatively evaluating the separate contributions of different levels of visual features, we found that the cortical organization of categories was explained by multiple levels of visual features but to different degrees. The spatial organization of biological objects was better explained by higher-level visual features (Fig. 8). This agreed with previous findings that higher layers in the CNN better explained the representational similarity of object categories in the inferior temporal cortex^18,19,72^. However, the spatial organizations of superordinate-level categories and background scenes were, surprisingly, best explained by mid-level visual features (Fig. 6 and Fig. 8). The difference in the best-explainable dimensions suggests that the cortical organizations of different categories are not uniquely driven by certain common dimensions. One possible interpretation of these findings is that the cortical organization of objects is attributable to various visual dimensions ranging from low to mid to high levels of visual features, and essentially to those dimensions that best characterize different objects. This organization principle also explains why the cortical representation of categories is partially, but not fully, explained by low-level features^60–62^, shapes^63,64^, or semantic information^59,65^. Importantly, the mid-level features, while much less known than low- and high-level features, largely explained the spatial organization of category representations.

## Materials and Methods

### Experimental data

We used and extended the human experimental data from our previous study^24^, according to experimental protocols approved by the Institutional Review Board at Purdue University with informed consent from all human subjects prior to their participation. Briefly, the data included the fMRI scans from three healthy subjects (Subject 1, 2, 3, all female) when watching natural videos. For each subject, the video-fMRI data were split into two independent datasets: one for training the encoding model and the other for testing it. For Subject 2 & 3, the training movie included 2.4 hours of videos; the testing movie included 40 minutes of videos; the training movie was repeated twice, and the testing movie was repeated ten times. For Subject 1, the training movie included not only those videos presented to Subject 2 and 3, but also 10.4 hours of new videos. The movie stimuli included a total of ~9,300 video clips manually selected from *YouTube*, covering a variety of real-life visual experiences. All video clips were concatenated in a random sequence and separated into 8-min sessions. Every subject watched each session of videos (field of view: 20.3^o^×20.3^o^) through a binocular goggle with the eyes fixating at a central cross (0.8^o^×0.8^o^). During each session, whole-brain fMRI scans were acquired with 3.5 mm isotropic resolution and 2 s repetition time in a 3-T MRI system. The volumetric fMRI data were preprocessed and co-registered onto a standard cortical surface template ^73^. More details about the movie stimuli, data preprocessing and acquisition are described elsewhere ^24^.

### Deep residual network

In line with previous studies^18–22,24,34^, a feedforward deep neural network (DNN) was used to model the cortical representations of natural visual stimuli. Here, we used a specific version of the DNN known as the deep residual network (ResNet), which had been pre-trained to categorize natural pictures with the state-of-the-art performance^32^. In the ResNet, 50 hidden layers of neuron-like computational units were stacked into a bottom-up hierarchy. The first layer encoded location and orientation-selective visual features, whereas the last layer encoded semantic features that supported categorization. The layers in between encoded increasingly complex features through 16 residual blocks; each block included three successive layers and a shortcut directly connecting the input of the block to the output of the block^32^. Compared to the DNNs in prior studies^19–21,24,34,72^, the ResNet was much deeper and defined more fine-grained hierarchical visual features. The ResNet could be used to extract feature representations from any input image or video frame by frame. Passing an image into the ResNet yielded an activation value at each unit. Passing a video yielded an activation time series at each unit as the fluctuating representation of a given visual feature in the video.

### Encoding models

For each subject, we trained an encoding model to predict each voxel’s fMRI response to any natural visual stimuli^74^, using a similar strategy as previously explored^20,22,24^. The voxel-wise encoding model included two parts: the first part was nonlinear, converting the visual input from pixel arrays into representations of hierarchical features through the ResNet; the second part was linear, projecting them onto each voxel’s fMRI response. The encoding model used the features from 18 hidden layers in the ResNet, including the first layer, the last layer, and the output layer for each of the 16 residual blocks. For video stimuli, the time series extracted by each unit was standardized (i.e. remove the mean and normalize the variance), and convolved with a canonical hemodynamic response function (HRF) with the peak response at 4s, and then down-sampled to match the sampling rate of fMRI.

The feature dimension was reduced by applying principle component analysis (PCA) first to each layer and then to all layers in ResNet. The principal components of each layer were a set of orthogonal vectors that explained >99% variance of the layer’s feature representations given the training movie. The layer-wise dimension reduction was expressed as equation (1).

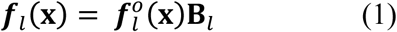

where 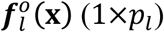 is the original feature representation from layer *l* given a visual input **x**, **B**_*l*_(*p*_*l*_×*q*_*l*_) consists of unitary columnar vectors that represented the principal components for layer *l*, *l*_*l*_(**x**) (1×*q*_*l*_) is the feature representation after reducing the dimension from *p*_*l*_ to *q*_*l*_.

Following the layer-wise dimension reduction, the feature representations from all layers were further reduced by using PCA to retain >99% variance across layers. The final dimension reduction was implemented as equation (2).

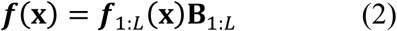

where 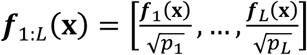 is the feature representation concatenated across *L* layers, **B**_1:*L*_ consists of unitary principal components of the layer-concatenated feature representations of the training movie, and **f**(**x**) (1×*k*) is the final dimension-reduced feature representation.

For the second part of the encoding model, a linear regression model was used to predict the fMRI response *r*_*v*_(**x**) at voxel *v* evoked by the stimulus **x** based on the dimension-reduced feature representation **f**(**x**) of the stimulus, as expressed by equation (3).

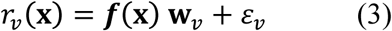

where **w**_*v*_ is a columnar vector of regression coefficients specific to voxel *v*, and ∈_*v*_ is the error term. As shown in equation (4), L_2_-regularized least-squares estimation was used to estimate **w**_*v*_ given the data during the training movie (individual frames were indexed by *i* = 1, ⋯, *N*), where the regularization parameter was determined based on nine-fold cross-validation.

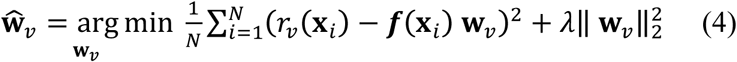

After the above training, the voxel-wise encoding models were evaluated for their ability to predict the cortical responses to the novel testing movie (not used for training). The prediction accuracy was quantified as the temporal correlation (*r*) between the predicted and observed fMRI responses at each voxel given the testing movie. Since the testing movie included five distinct sessions, the prediction accuracy was evaluated separately for each session, and then averaged across sessions. The significance of the voxel-wise prediction accuracy was evaluated with a block-permutation test^75^ (corrected at false discovery rate (FDR) *q* < 0.01), as used in our prior study^24^.

We also evaluated the correspondence between the hierarchical layers in ResNet and the hierarchical cortical areas underlying different stages of visual processing, in line with previous studies^18–24,34^. For this purpose, we calculated the variance of the response at a voxel explained by the visual features in single layers. Specifically, the features extracted from the testing movie were kept only for one layer in the ResNet, while setting to zeros for all other layers. Through the voxel-wise encoding model, the variance (measured by R-squared) of the response explained by the single layer was calculated. For each voxel, we identified the best corresponding layer with the maximum explained variance and assigned its layer index to this voxel. The assigned layer index indicated the processing stage this voxel belonged to.

We also tested whether the deeper ResNet outperformed the shallower AlexNet^33^ in predicting cortical responses to natural movies, taking the latter as the benchmark given its state-of-the-art encoding performance in prior studies^20,21,24^. For this purpose, we trained and tested similar encoding models based on the AlexNet with the same analysis of the same dataset. We compared the prediction accuracy between ResNet and AlexNet for regions of interest (ROIs) defined in an existing cortical parcellation^76^, and further evaluated the statistical significance of their difference using a paired t-test (p<0.001) across all voxels within each ROI. Considering the noise in the data, we also calculated the noise ceiling of the predictability at each voxel. The noise ceiling indicated the maximum accuracy that a model could be expected to achieve given the level of noise in the testing data^77^. The noise and signal in fMRI were assumed to follow Gaussian distribution and the mean of noise was zero. For each testing session, we estimated the noise level and the mean/SD of the signal for every voxel. We used Monte Carlo simulation to obtain the noise ceiling. For each simulation, we generated a signal from the signal distribution, and generated a noisy data by adding the signal and the noise drawn from the noise distribution, and calculated the correlation between the signal and the data. We performed 1,000 simulations for each testing session, and took the median correlation as the noise ceiling. The ceiling was then averaged across sessions.

### Human-face representations with encoding models and functional localizer

The ResNet-based encoding models were further used to simulate cortical representations of human faces, in comparison with the results obtained with a functional localizer applied to the same subjects. To simulate the cortical “face” representation, 2,000 human-face pictures were obtained by Google Image search. Each of these pictures was input to the voxel-wise encoding model, simulating a cortical response map as if it were generated when the subject was actually viewing the picture, as initially explored in previous studies^22,24^. The simulated response maps were averaged across all the face pictures, synthesizing the cortical representation of human face as an object category.

To validate the model-synthesized “face” representation, a functional localizer^78^ was used to experimentally map the cortical face areas on the same subjects. Each subject participated in three sessions of fMRI with a randomized block-design paradigm. The paradigm included alternating ON-OFF blocks with 12s per block. During each ON block, 15 pictures (12 novel and 3 repeated) from one of the three categories (face, object, and place) were shown for 0.5s per each picture with a 0.3s interval. The ON blocks were randomized and counter-balanced across the three categories. Following the same preprocessing as for the video-fMRI data, the block-design fMRI data were analyzed with a general linear model (GLM) with three predictors, i.e. face, object, and place. Cortical “face” areas were localized by testing the significance of a contrast (face>object and face > place) with p<0.05 and Bonferroni correction.

### Synthesizing cortical representations of different categories

Beyond the proof of concept with human faces, the similar strategy was also extended to simulate the cortical representations of 80 categories through the ResNet-based encoding models. The category labels were shown in Fig. 3. These categories were mostly covered by the video clips used for training the encoding models. For each category, 800 pictures were obtained by Google Image search with the corresponding label, and were visually inspected to replace any exemplar that belonged to more than one category. The cortical representation of each category was generated by averaging the model-simulated response map given every exemplar within the category.

### Category selectivity

Following the above analysis, cortical representations were compared across categories to quantify the category selectivity of various locations and ROIs. For each voxel, its selectivity to category *i* against other categories *i*^c^ was quantified with equation (5), as previously suggested^36^.

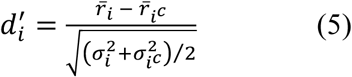

where 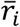 and 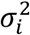 are the mean and variance of the responses to the exemplars in category *i*, and 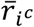 and 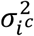 were counterparts to all exemplars in other categories *i*^c^. Irrespective of any specific category, the general category-selectivity for each voxel was its maximal *d*′ index among all categories, i.e. 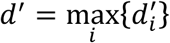. A *d*′ index of zero suggests non-selectivity to any category, and a higher *d*′ index suggests higher category-selectivity. The category selectivity of any given voxel was also inspected by listing the categories in a descending order of their representations at the voxel. We also obtained the ROI-level category selectivity by averaging the voxel-wise selectivity across voxels and subjects. ROIs were defined in an existing cortical parcellation^76^.

### Categorical similarity and modularity in cortical representation

To reveal how the brain organizes categorical information, we assessed the similarity (i.e. spatial correlation across the predictable voxels with q<0.01 in permutation test and prediction accuracy r>0.2) in cortical representations between categories. Based on such inter-category similarity, individual categories were grouped into clusters using k-means clustering ^79^. The goodness of clustering was measured as the modularity index, which quantified the inter-category similarities within the clusters relative to those regardless of the clusters^80^.

The similarity in cortical representation between different categories was compared with their similarity in semantic meaning. The semantic similarity between categories was evaluated as the Leacock-Chodorow similarity^37^ between the corresponding labels based on their relationships defined in the WordNet^81^ – a directed graph of words (as the nodes) and their *is-a* relationships (as the edges). The correlation between the cortical and semantic similarities was evaluated across all pairs of categories.

### Layer-wise contribution to cortical categorical representation

We also asked which levels of visual information contributed to the modular organization of categorical representations in the brain. To answer this question, the cortical representation of each category was dissected into multiple levels of representations, each of which was attributed to one single layer of features. For a given category, the features extracted from every exemplar of this category were kept only for one layer in the ResNet, while setting to zeros for all other layers. Through the voxel-wise encoding model, the single-layer visual features were projected onto a cortical map that only represented a certain level of visual information shared in the given category. The similarity and modularity in cortical representations of individual categories were then re-evaluated as a function of the layer in the ResNet. The layer with the highest modularity index contributed the most to the modular organization in cortical categorical representation. The features encoded by this layer were visualized for more intuitive understanding of the types of visual information underlying the modular organization. The feature visualization was based on an optimization-based technique^82^. Briefly, to visualize the feature encoded by a single unit in the ResNet, the input to the ResNet was optimized to iteratively maximize the output from this unit, starting from a Gaussian random pattern. Four optimized visualizations were obtained given different random initialization.

### Categorical representation in nested hierarchical spatial scales

Considering object categories were defined hierarchically in semantics^81^, we asked whether there were spatial and representational hierarchies underlying the hierarchy of categorization^3^. More specifically, we tested whether the representational similarity and distinction in a larger spatial scale gave rise to a coarser level of categorization, whereas the representation in a smaller spatial scale gave rise to a finer level of categorization. To do so, we first examined the category representation in the scale of the entire visual cortex predictable by the encoding models, and clustered the categories into multiple modules by using the modularity analysis of the representational similarity in this large scale. The resulting modules of categories were compared with the superordinate-level semantic categories. Then, we focused on a finer spatial scale specific to the regions where category representations overlapped within each module in contrast to 50,000 random and non-selective objects (p<0.01, two-sample t-test, Bonferroni correction). Given the spatial similarity of category representation in this finer scale, we defined sub-modules within each module using the same modularity analysis as for the large-scale representation. The sub-modules of categories were compared and interpreted against semantic categories in a finer level.

## Data Availability

The datasets generated during and/or analyzed during the current study are available from the corresponding author on reasonable request.

### >Acknowledgement

The authors are thankful to Dr. Xiaohong Zhu and Dr. Byeong-Yeul Lee for constructive discussion, and Kuan Han for his assistance in collecting natural images, and Yizhen Zhang for her help in acquiring fMRI. The research was supported by NIH R01MH104402, MH111413, and Purdue University.

## Authors’ Contributions

H. Wen, J. Shi, W. Chen and Z. Liu designed the study. H. Wen and J. Shi collected the data. H. Wen analyzed the data. H. Wen collected the natural images. H. Wen, Z. Liu and W. Chen wrote the paper. All authors contributed to data interpretation.

## Additional Information

### Competing financial interests

The authors have no competing interests as defined by Nature Publishing Group, or other interests that might be perceived to influence the results and/or discussion reported in this paper.

